# The cell wall lipoprotein CD1687 acts as a DNA binding protein during deoxycholate-induced biofilm formation in *Clostridioides difficile*

**DOI:** 10.1101/2022.11.29.518320

**Authors:** Emile Auria, Lise Hunault, Patrick England, Marc Monot, Juliana Pipoli Da Fonseca, Mariette Matondo, Magalie Duchateau, Yannick D.N. Tremblay, Bruno Dupuy

## Abstract

The ability of bacterial pathogens to establish recurrent and persistent infections is frequently associated with their ability to form biofilms. *Clostridioides difficile* infections have a high rate of recurrence and relapses and it is hypothesised that biofilms are involved in its pathogenicity and persistence. Biofilm formation by *C. difficile* is still poorly understood. It has been shown that specific molecules such as deoxycholate (DCA) or metronidazole induce biofilm formation, but the mechanisms involved remain elusive. In this study, we describe the role of the *C. difficile* lipoprotein CD1687 during DCA-induced biofilm formation. We showed that the expression of *CD1687*, which is part of an operon within the *CD1685-CD1689* gene cluster, is controlled by multiple transcription starting sites and some are induced in response to DCA. Only CD1687 is required for biofilm formation and the overexpression of CD1687 is sufficient to induce biofilm formation. Using RNAseq analysis, we showed that CD1687 affects the expression of transporters and metabolic pathways and we identified several potential binding partners by pull-down assay, including transport-associated extracellular proteins. We then demonstrated that CD1687 is surface exposed in *C. difficile*, and that this localization is required for DCA-induced biofilm formation. Given this localization and the fact that *C. difficile* forms eDNA-rich biofilms, we confirmed that CD1687 binds DNA in a non-specific manner. We thus hypothesize that CD1687 is a component of the downstream response to DCA leading to biofilm formation by promoting interaction between the cells and the biofilm matrix by binding eDNA.

## Introduction

Gastrointestinal infections are a major public health issue. In high-income countries, the Gram-positive spore forming anaerobe *Clostridioides difficile* is the leading cause of nosocomial diarrhea and colitis in adults receiving antibiotic treatments (1,2). Moreover, *C. difficile* infections (CDI) can be persistent, which is a major challenge in the management of CDI following anti-*C. difficile* antibiotic treatment. Recurrent CDI occur in more than 20% of patients that receive antibiotics to treat their first CDI episode and this rate increases following new episodes (3,4). The causes of recurrences have not been fully elucidated. Recurrence can be caused by either reinfection with a new strain or relapse with the same strain, suggesting that *C. difficile* can persist in the gastrointestinal tract (5). Relapses were initially correlated with *C. difficile* ability to sporulate during the infection and resist antibiotic treatment (6,7). However, relapses are also hypothesized to be associated with the persistence of *C. difficile* as a biofilm (8,9). Persistent and chronic infections caused by different pathogens are known to be associated with biofilm formation (10). It is estimated that at least 60% of all nosocomial and chronic bacterial infections are biofilm-associated (11). In support of this hypothesis, *C. difficile* was recently showed to integrate biofilms formed by the colonic microbiota and this biofilm acted as a reservoir for persistence and recurrence in a laboratory model of CDI (9).

Biofilms are structured communities of microorganisms associated with surfaces and encased in a self-produced extracellular matrix, which varies between bacterial species (12). *C. difficile* can form biofilms as a single species or with other bacteria on various abiotic surfaces and several *in vitro* systems (9,13–15). Moreover, *C. difficile* can integrate *in vivo* multi-species communities during a mouse infection, suggesting its ability to integrate mucosal biofilms (16). Additionally, *C. difficile* can form patchy glycan-rich biofilm-like structures in a mono-associated mouse model (17). Although *C. difficile* can integrate multispecies biofilms in the gastrointestinal tract, there is limited knowledge on the biology of *C. difficile* biofilm formation in response to the gastrointestinal environment. During an infection, pathogens encounter several environmental factors including the presence of antibiotics, bile salts, osmotic pressure and varying nutrient sources and these are known to be important signals for biofilm formation during colonization (18,19). Interestingly, *C. difficile* would face different challenges during dysbiosis as it changes of the nutritional environment, bile salt metabolism, and osmotic and oxidative/nitrosative stresses, (20). Any of these factors could induce biofilm formation. For example, sub-inhibitory concentrations of antibiotics used to treat CDI enhances biofilm formation *in vitro* (21,22). Furthermore, we recently demonstrated that sub-inhibitory concentrations of the secondary bile salt deoxycholate (DCA) enhances *C. difficile* biofilm formation (15). In the DCA-induced biofilm, vegetative cells are protected from the toxicity of DCA as well as antibiotics and antimicrobial peptides (15). We showed that biofilms induced by DCA are formed due to metabolic adaptation and reprogramming that are dependent on the available nutrients and excreted metabolites. Overall, excreted pyruvate is critical for the induction of biofilm formation (23).

In addition to environmental factors inducing biofilm formation, several cellular factors, including cell surface components and regulators, have been shown to influence biofilm formation by *C. difficile* (24). Among the genes that were up-regulated in response to DCA, a gene encoding a lipoprotein (CD1687) is essential for biofilm formation in response to DCA (15). The aim of this study was to characterize the role of CD1687 during biofilm formation by *C. difficile* in response to DCA. We demonstrated that CD1687 is exposed and active at the surface of the bacteria and that it binds DNA *in vitro*. This suggests that CD1687 acts as a protein anchoring the cells to the extracellular DNA (eDNA) present in the biofilm matrix.

## Methods

### Bacterial strains and culture conditions

Bacterial strains and plasmids used in this study are listed in Table S1 *C. difficile* strains were grown anaerobically (5% H2, 5% CO2, 90% N2) in TY medium (30g/L tryptone, 20g/L yeast extract) or in BHISG medium (BHI with 0.5% (w/v) yeast extract, 0.01 mg/mL cysteine and 100mM glucose) and supplemented with cefoxitin (250μg/ml), D-cycloserine (8μg/ml) and thiamphenicol (15 μg/ml) when necessary. Additionally, 100ng/mL of anhydrotetracycline (ATC) was added to induce the *P*_*tet*_ promoter of pRPF185 vector derivatives in *C. difficile. E. coli* strains were grown in LB broth supplemented with chloramphenicol (15μg/mL) and ampicillin (100μg/mL).

### Biofilm assays

Overnight cultures of *C. difficile* grown in TY medium with appropriate antibiotics were diluted to 1/100 into fresh BHISG containing the desired supplements (240μM DOC, 100ng/mL ATC or both). Depending on the assay, the diluted cultures were then aliquoted either with 1mL per well in 24-well plates (polystyrene tissue culture-treated plates, Costar, USA) or with 200μL in 96-well plates (polystyrene black tissue-culture-treated plates, Greiner Bio One, Austria). The plates were incubated at 37°C in an anaerobic environment for 48h. Biofilm biomass was measured in the 24-well plates using an established method (15). For biofilm assays in 96-well-plates used for microscopy, spent medium was carefully removed by pipetting and 200μL PBS supplemented with 4% of paraformaldehyde (PFA) were added. Plates were incubated for an hour at room temperature and the media was then carefully removed by pipetting before adding PBS for 48h at 4°C. In all assays, sterile medium was used as a negative control and a blank for the assays.

### Gene deletion in *C. difficile*

Gene deletion in *C. difficile* was performed as described in Peltier *et al*. (2020) (25). Regions upstream and downstream of the genes of interest were PCR-amplified using primer pairs described in Table S1. PCR fragments and linearized pDIA6754 (25) were mixed and assembled using Gibson Assembly (NEB, France) and transformed by heat shock in *E. coli* NEB 10β strain. The plasmid constructions were verified by sequencing and plasmids with the right sequences were transformed in *E. coli* HB101 (RP4). The resulting strains were used as donors in a conjugation assay with the relevant *C. difficile* strains. Deletion mutants were then obtained using a counter-selection as described in Peltier et al (2020) (25).

### Protein extraction from *C. difficile* and pull-down assay

*C. difficile* strains were anaerobically grown for 48h in 20mL BHISG cultures with ATC in tubes. Cells and biofilms were harvested by centrifugation (10 min; 13000 rpm; 4°C) and washed in a cold phosphate buffer (50mM; pH=7.0; 4°C). Cells were then resuspended in 1 ml of the same phosphate buffer containing the purified catalytic domain of the endolysin CD27L (3μg/mL) and suspension was incubated 1h at 37°C to lyse the bacterial cells. The pull-down assay was then performed with Ni-NTA beads as described in supplementary data for CD1687 purification from *E. coli* expression.

### RNA isolation, qRT PCR

Cells were grown in 24-well plates and 10 wells per plate were used to produce one replicate for one condition. For biofilm conditions, the supernatant was removed by inverting the plate and the biofilms were carefully washed twice then resuspended in 3 mL of PBS. In other conditions, the whole bacterial population was collected and cells were harvested by centrifugation (10 min, 5000 rpm, 4°C) and resuspended in 1 ml of PBS. Cell suspensions in PBS were finally centrifuged (10 min, 5000 rpm, 4°C) and the pellets were frozen at -80°C until further use. Extraction of total RNA from the bacteria and qRT PCR assay were performed as described in Saujet *et al* (2011) (26).

### Whole transcriptome sequencing and analysis

Transcriptome analysis for each condition was performed using 4 independent RNA preparations. Libraries were constructed using the Illumina Stranded Total RNA Prep Ligation with RiboZero Plus (Illumina, USA) kit. The ribodepletion step was carried using specific probes synthesized specifically to target *C. difficile* ribosomal sequences. After ribodepletion, libraries were prepared according to the supplier’s recommendations. RNA sequencing was performed on the Illumina NextSeq 2000 platform using 67 bases for a target of 10M reads per sample.

### Electromobility shift assays (EMSA)

Only freshly purified CD1687 from *E. coli* were used in these assays. CD1687 (from 0.5μM to 16μM) was incubated with DNA (pUC9 or PCR product) in 10μl of sodium phosphate buffer (50mM; pH=8.0) for 30 min at room temperature. Samples were loaded and migrated on TAE buffered agarose gels (1% w/v) for 90min at 100V. Controls were performed with CD1687 denatured at 100°C for 15 min before the assay. Gels were stained with ethidium bromide and pictures were taken with an Amersham ImageQuant 800 (Cytiva). The pUC9 plasmid was prepared from *E. coli* stock using the Nucleospin plasmid kit (Macherey-Nagel, Germany) and the PCR amplicon used was generated using *C. difficile* 630*Δerm* as the DNA template and primers targeting the region of *CD1438* (Table S1). gDNA was extracted from cell culture using the DNeasy Blood & Tissue Kit (QIAGEN, Netherlands).

### 5’RACE experiment

A 5’RACE was performed using the 5’ RACE System for Rapid Amplification of cDNA Ends, version 2.0 kit (Invitrogen, USA). Briefly, cDNA was generated by reverse transcription from total RNA extract followed by degradation of the RNA. dC-tailing was then performed with the cDNA and the resulting dC-tailed DNA was used as the template in PCR as described in the kit instructions. The PCR products were analyzed by agarose gel electrophoresis (1% agarose in TAE buffer). To identify the transcription start sites, PCR products were inserted into the pGEM-T easy vector kit as described by the manufacturer (Promega, USA). Insert were then PCR-amplified and the resulting PCR products were sequenced.

### Epifluorescence microscopy

For microscopy, 48h biofilms were generated in 96-well plates (black, Greiner) as described above, washed and 50μl of the polyclonal anti-CD1687 antibodies diluted in PBS (400 ng/mL) was then added to each well and incubated overnight at 4°C. The wells were carefully washed twice with PBS followed by the addition of a solution containing DAPI (1/1000 dilution) and secondary antibodies (goat anti-rabbit conjugated with Texas Red; 1/5000 dilution; Invitrogen) in PBS. The plates were incubated at room temperature for 2h. Wells were then carefully washed with PBS and 200μl of fresh PBS was added for data acquisition. Images were taken with the Nikon Eclipse Ti inverted microscope (Nikon, Japan).

### Statistical analysis

The biofilm assays and RT-qPCR were analysed using a one-way ANOVA test followed by either a Tukey’s multiple comparison test or a Dunnett’s multiple comparison test.

## Data Availability

RNA-Seq data generated in this study are available in the NCBI-GEO with the accession number GSE218475. The mass spectrometry proteomics data have been deposited to the ProteomeXchange Consortium via the PRIDE partner repository with the dataset identifier PXD038282.

## Results

### Genes of the *CD1685-CD1689* locus form an operon but multiple transcription start site control their expression

In previous transcriptomic experiments, we observed that the majority of genes in the *CD1685 - CD1689* cluster were up-regulated in the 48h DCA-induced biofilm formed by *C. difficile* strain 630Δ*erm* (15,23). However, inactivation of CD1687 but not CD1688 prevented DCA-induced biofilm formation. To verify that the *CD1685-CD1689* genes formed an operon, RT-PCR experiments were performed with RNA extracted from cells grown under biofilm inducing conditions (BHISG with 240μM DCA). We observed a unique transcript spanning *CD1685* to *CD1689* suggesting the presence of at least one polycistronic mRNA at this locus (Fig 1a). We then performed qRT-PCR to confirm that the five genes were up-regulated at 48h in the presence of DCA and only small difference in the fold changes were seen (Fig S1a).

**Figure 1:**
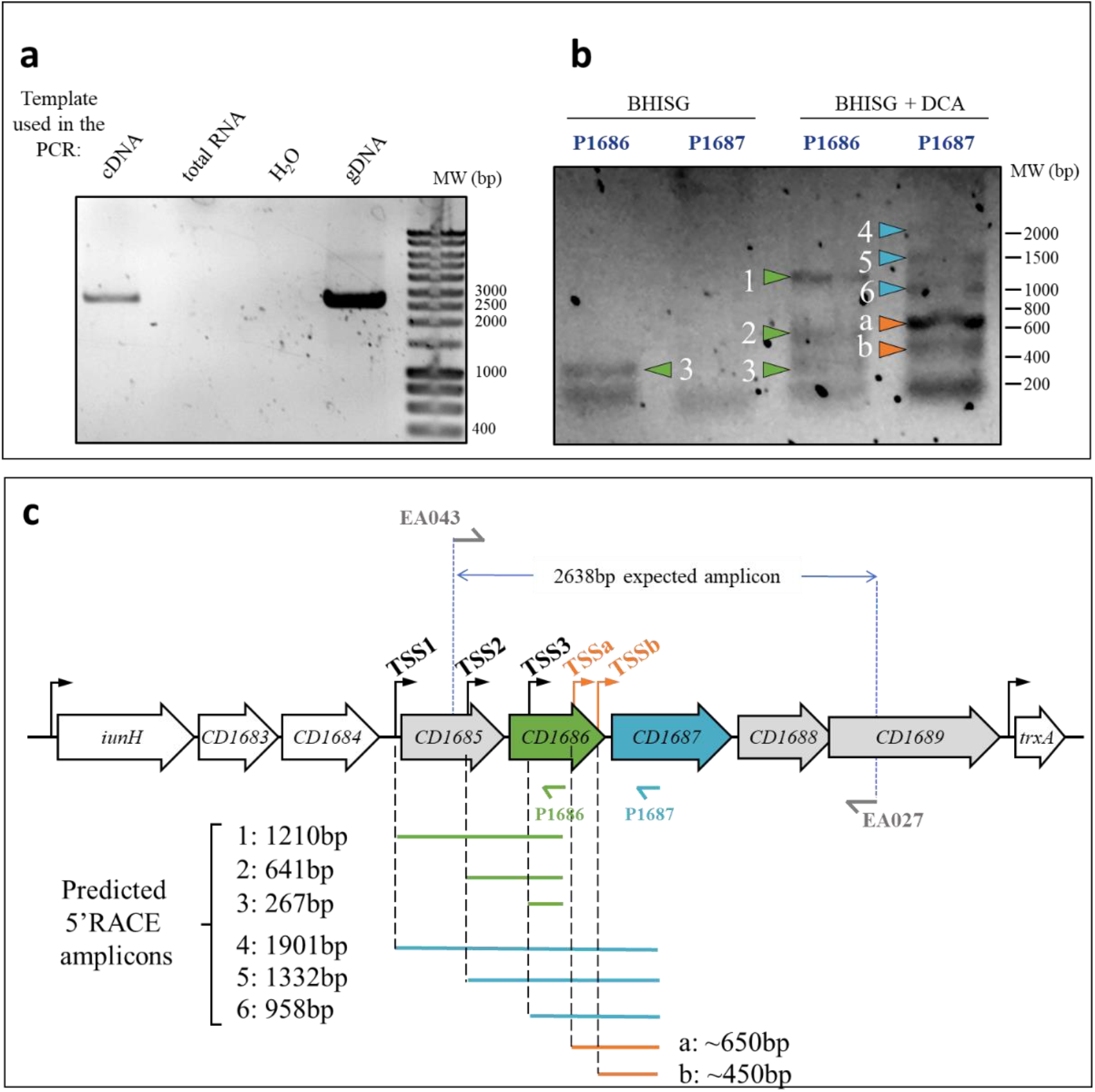
The *CD1685-CD1689* cluster in *C. difficile* strain 630Δ*erm* forms an operon with multiple transcription start sites. **a**. RT-PCR performed with primers EA043 and EA027 (Table S1) from various nucleic acid templates. cDNA was obtained using the EA027 primer with total RNA extracted from 48h biofilms grown in BHISG supplemented with DCA (240μM). **b**. 5’RACE results from amplification of the poly-guanylated cDNA obtained respectively with the EA021 and EA018 primers (Table S1), then the P1686 or P1687 primers along with the universal amplification primer (AAP) from the 5’RACE kit. The RNA was extracted from 48h cell cultures grown under biofilm inducing conditions (BHISG + 240μM DCA) or non-biofilm-inducing conditions (BHISG). **c**. Organisation of the *CD1685-CD1689* cluster, the location of the primers used for RT-PCR and the amplicons from the 5’RACE results using the P1686 or P1687 primers (amplicon sizes were predicted from the TSS identified by Soutourina *et al*. (2020) and Fuchs *et al*. (2021). TSS: Transcriptional Start Site; cDNA: complementary DNA; gDNA: genomic DNA.

When looking at our previous RNAseq experiments, we observed a mapping bias of the sequencing reads favouring CD1687, CD1688 and CD1689 (Fig S1b). Interestingly, recent analyses predicted three transcription starting sites (TSS) for the CD1685-CD1689 locus: one upstream of the *CD1685* gene (TSS1), one upstream the *CD1686* gene (TSS2) and one in the coding sequence of *CD1686* (TSS3) (27,28) (Fig 1c). To confirm the existence of multiple TSS, 5’-RACE experiments were performed with total RNA extracted from cells grown for 48h in BHISG with DCA (i.e. biofilm-inducing) or without DCA (i.e. non-biofilm inducing). The initial reverse transcriptions were performed with two primers annealing either the coding sequence of *CD1686* (P1686) or the coding sequence of *CD1687* (P1687) (Fig 1bc). In the absence of DCA, only one amplicon was observed, which is associated with the TSS inside *CD1686*. This amplicon was detectable when the P1686 primer was used but not with the P1687 primer. In the presence of DCA, we observed amplicons corresponding to the three predicted TSS with either primer (P1686 or P1687) and two additional amplicons were detected with P1687. This suggest that these two additional TSS (TSSa and TSSb; Fig 1c) are active in the presence of DCA and one of these (TSSa) appears to be the most active of all TSS (Fig1b). Each amplicon was sequenced (Table S2) and the location of TSS1, TSS2 and TSS3 closely matched their predicted location. However, high variation of the sequences for TSSa and TSSb made it difficult to identify their exact location. Overall, the transcription of the *CD1685-CD1689* operon is initiated from multiple TSS in the presence of DCA, suggesting that multiple factors are integrated to regulate the expression of the CD1685-CD1689 operon to reflect the state of the bacterial population.

### Overexpressing CD1687 induces biofilm formation in the absence of DCA

We previously inactivated *CD1687* using the Clostron system (15) but this approach is known to have some limitations. To confirm that only *CD1687* was required for biofilm formation, deletion of *CD1686, CD1687* and *CD1688-CD1689* were generated (Fig S2a). As observed before, only the deletion of *CD1687* negatively affected biofilm-formation and complementation restore the phenotype (Fig S2bc). Interestingly, deletion of *CD1686* removed TSS3, TSSa and TSSb suggesting that TSS1 and/or TSS2 are sufficient for the transcription of CD1687 in the presence of DCA resulting in biofilm formation.

Since CD1687 is required for DCA-induced biofilm formation and previously localised in the cell wall fraction (15), we hypothesized that CD1687 is a DCA-sensing protein. To test this hypothesis, we verified the ability of CD1687 to directly interact with DCA using surface plasmon resonance. We showed that CD1687 can interact with DCA (Fig S3). However, the dissociation constant is high (Kd of 1.65±0.58mM), and the estimated stoichiometry of the interaction is of 5±1 DCA molecules for one CD1687 protein, which implies that the interaction is not specific.

Interestingly, we observed an increase in biofilm formation in the presence and, to a certain extent, in absence of DCA when the Δ*1687* mutant was complemented with an inducible plasmid-borne CD1687 (pDIA6920) (Fig S2C). Although the increase was not significant, it suggested that CD1687 could induce biofilm formation in the absence of DCA. To test this hypothesis, pDIA6920 was introduced in the wild type strain and its ability to form biofilm in the absence of DCA was evaluated with and without the addition of the inducer ATC. When CD1687 was overexpressed, a stronger biofilm was detectable at 24h and 48h (Fig 2). Taken together, our results suggest that CD1687 expression is critical for biofilm formation which does not require DCA for its activity.

**Figure 2:**
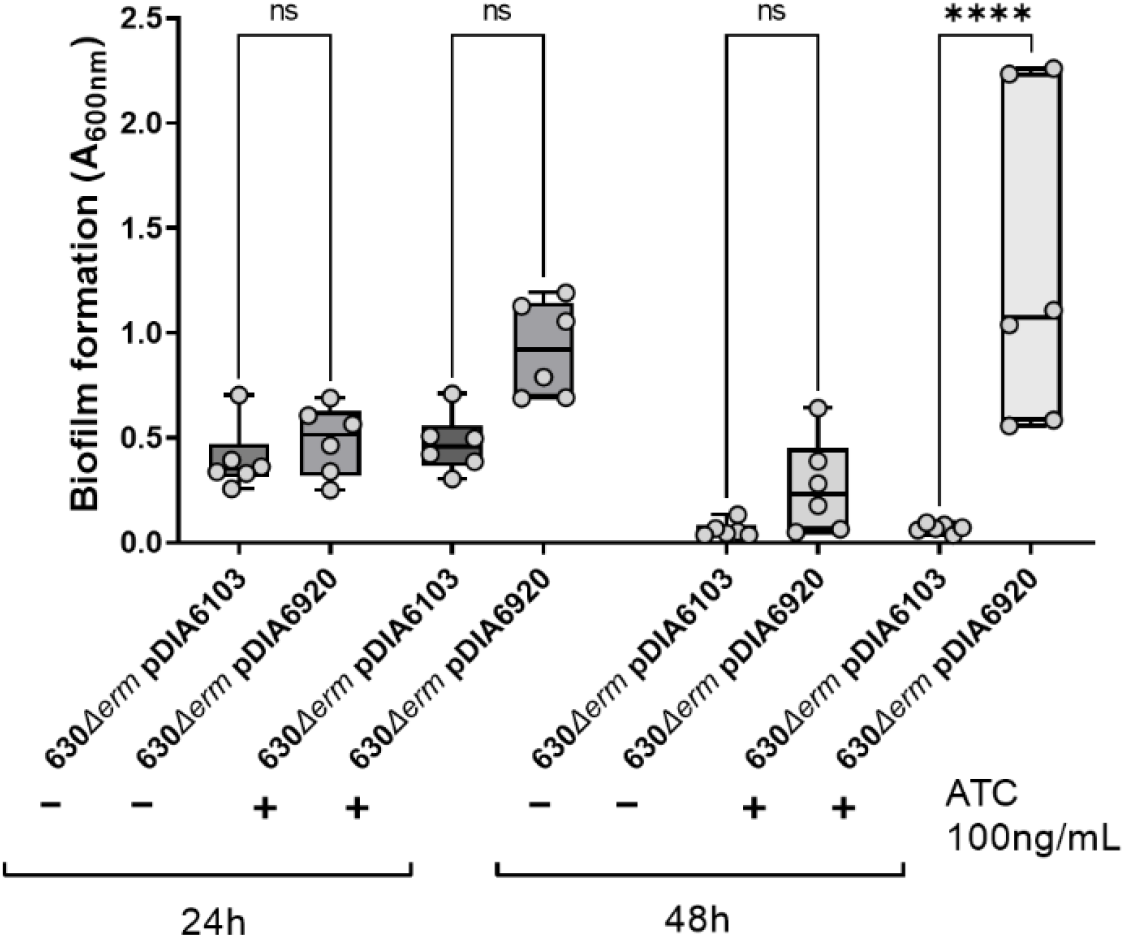
Overexpression of CD1687 induces biofilm formation in the absence of DCA. Biofilms formation was assayed 24h or 48h after inoculation in BHISG +/-ATC (100ng/mL) with the wild type strain (630Δ*erm*) containing either a control empty vector (pDIA6103) or the vector allowing the expression of CD1687 under the inducible *P*_*tet*_ promoter (pDIA6920). Each data point represents an independent biological replicate composed of 2 to 4 technical replicates. Asterisks indicate statistical significance with a one-way ANOVA test followed by a Tukey’s multiple comparison test (ns: not significant; ****: p<0.0001).

### CD1687 affects the expression of several transporter and metabolic priorities

As CD1687 is essential for DCA-induced biofilm formation and its overexpression can induce biofilm formation in the absence of DCA, we sought to identify genes controlled by CD1687 during the biofilm formation process. To do so, we performed two transcriptomic analyses: one comparing the wild type and the *Δ1687* mutant grown in presence of DCA for 24h, and the second comparing the wild-type grown in absence of DCA for 24h overexpressing CD1687 or not overexpressing CD1687.

A total of 527 genes had a significant differential expression with a fold change <0.5 or >2 in the wild type strain compared to the *Δ1687* mutant under biofilm inducing conditions (+DCA) (Fig 3). In the presence of DCA, CD1687 seems to mainly downregulate the cell wall reticulation (*vanY2Y3*) as well as several uncharacterized regulators (Fig S4, Table S3). There seems to be a shift in membrane transporters that may result in an increase in the importation of branched-chain amino acids, iron and a change in sugar transport (Table S3). In terms of metabolism, the cells shift from the utilization of succinate (*CD2338-CD2344*), the Wood-Ljungdahl pathway and the biosynthesis of aromatic amino acids to the fermentation of acetoin, leucine, branched chain amino acids and glycine (Fig S4, Table S3).

**Figure 3:**
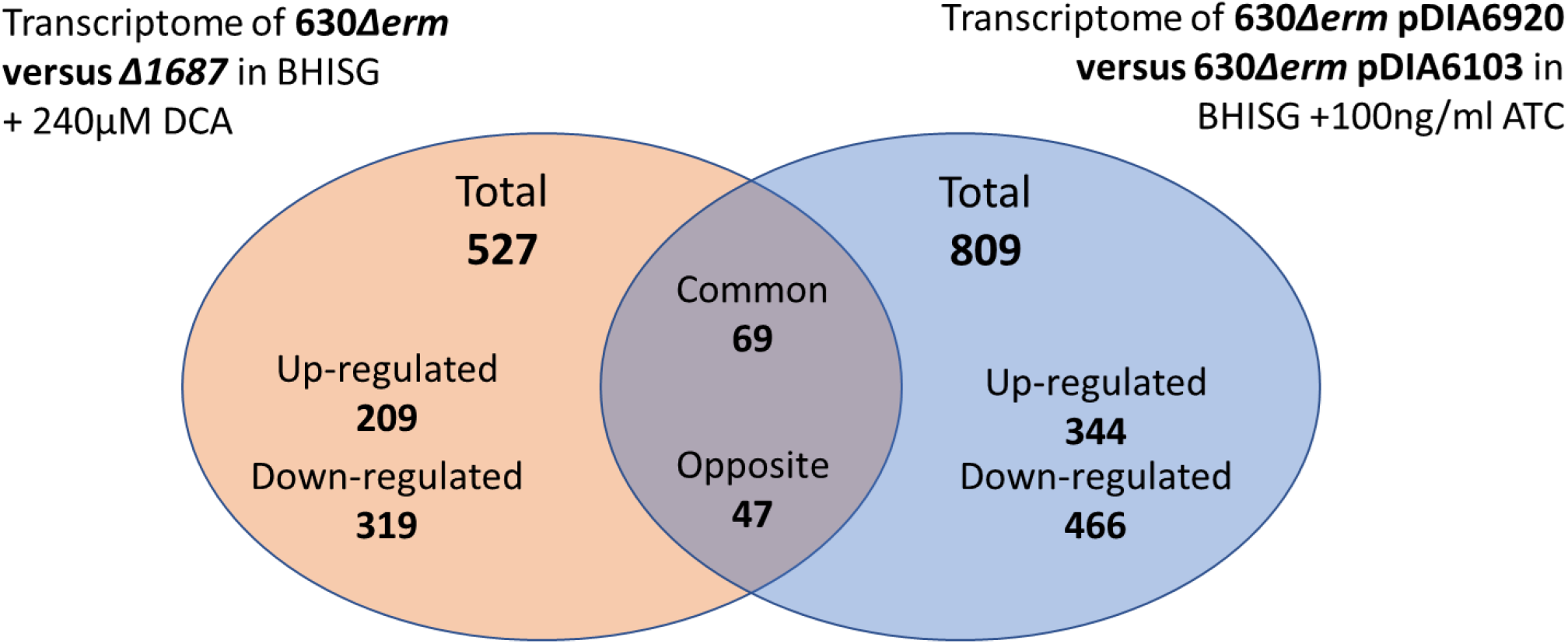
Differences in gene expression in the two transcriptomics experiments. Venn diagram of the genes differentially regulated in the two transcriptomics experiments performed in this study (Table S4).

When CD1687 was overexpressed, 809 genes were differentially expressed, 343 genes were up regulated and 466 were down-regulated (Fig 3). As described in Fig S4, changes in gene expression indicate a shift in transporters, metabolism and regulation. Specifically, the expression of several sugar transporters is increased whereas the expression of the branched chain amino acids, methionine, alanine and glycine transporters is down-regulated (Table S3). In terms of metabolism, genes involved in acetoin utilization, Stickland fermentations involving aromatic amino acids or leucine, the Wood-Ljungdahl pathway and the pentose phosphate pathway are up-regulated as well as those involved in the biosynthesis of several amino acids such as histidine, isoleucine, valine and cysteine (Table S3). The *dltABCD* operon is up-regulated suggesting an increase of the D-alanylation of the teichoic acids (*dltABCD*). Interestingly, we noted that the gene cluster encoding the flagellum and genes associated with sporulation were up-regulated.

When we compared both transcriptomic analyses, few genes overlapped between both analyses. Only 69 genes changed in the same direction whereas 47 genes were regulated in opposite direction (Fig 3). The remaining 1220 genes were differentially expressed only under either condition (Fig 3). The genes that were regulated in both conditions include those involved in cysteine synthesis (*cysE, cysK*), leucine utilization in Stickland fermentation (*hadABCI*), acetoin fermentation (*acoABCL*), cell wall proteins (*cwp9, cwp12*), some transporters (*alsT* transporting alanine or glycine, *rbsK* transporting ribose) and regulation (*sinRR’*). Overall, this suggests that CD1687 induces metabolic reorganization, including those occurring in response to DCA that leads to biofilm formation (23).

However, these changes do not fully align with our previous analyses (23). We previously observed that DCA causes the up-regulation of gene involved in butanoate, lactate and acetate fermentations, a shift in Stickland fermentations from the use of aromatic amino acids to the use of branched chain amino acids and glycine, and the down-regulation of genes involved in glycolysis, glucose intake and sporulation (23). These changes were not observed when CD1687 was overexpressed suggesting that CD1687 is not involved in those processes or does not mediate the immediate response to DCA. CD1687 is probably part of the downstream response and may interact with other proteins to promote these changes.

### CD1687 interacts with several cell wall proteins

Given that CD1687 is a cell wall protein (15) and does not have a transmembrane domain, we hypothesized that CD1687 induces transcriptional changes by transmitting external signals by interacting with membrane proteins. To find these potential proteins, we performed a pull-down assay using crude extracts of *C. difficile* cells overexpressing a hexahistidine tagged CD1687 in BHISG without DCA, and 43 proteins were captured (Table S5). Among the identified proteins, four are predicted to be membrane proteins and include a component of sugar transporter (CD2667) and a sodium symporter (CD2693). We also identified four proteins that belong to the large family of solute binding proteins associated with ABC transporters and one nucleotide phosphodiesterase (CD0689). These five proteins could be involved in signal transport and cellular response leading adaptation in different environmental conditions (29,30). Among the membrane proteins, we also found a putative lipoprotein (CD0747) and a LCP (LytR-CpsA-Psr) family protein (CD2766) involved in the cell wall polysaccharide assembly (31). We noted that only one encoding gene of protein partners (CD0037) was up-regulated in both transcriptomes (Table S5), which is typically localized in the cytoplasm. Since most of the membrane proteins identified by the pull-down experiment are cell wall proteins involved in membrane transport, it is possible that CD1687 directly affect transport of different nutrients and is consistent with the observed effect in our transcriptomes.

### CD1687 is exposed at the cell surface

Since CD1687 was detected in the cell wall fraction (15), we wondered whether CD1687 is exposed at the cell surface. To verify this, we performed epifluorescence microscopy analysis of *C. difficile* 630Δ*erm* strain and its derivatives using rabbit polyclonal antibodies raised against CD1687. When grown 48 hours in BHISG with or without DCA, no signal was observed in the Δ*1687* mutant confirming the specificity of our antibody (Fig 4 and Fig S5). For the wild-type strain, we observed a weak signal when grown in absence of DCA, confirming that this protein is expressed at low levels under non-biofilm inducing conditions. In the presence of DCA, the signal was stronger in the presence of DCA, although the expression of CD1687 was not homogeneous in the population. In contrast, the signal for CD1687 is homogenous in the population of the complemented Δ*1687* strain (Fig 4 and Fig S5). Since the cells were not permeabilized during the experiment, we concluded that CD1687 is exported to the cell wall and exposed at the cell surface.

**Figure 4:**
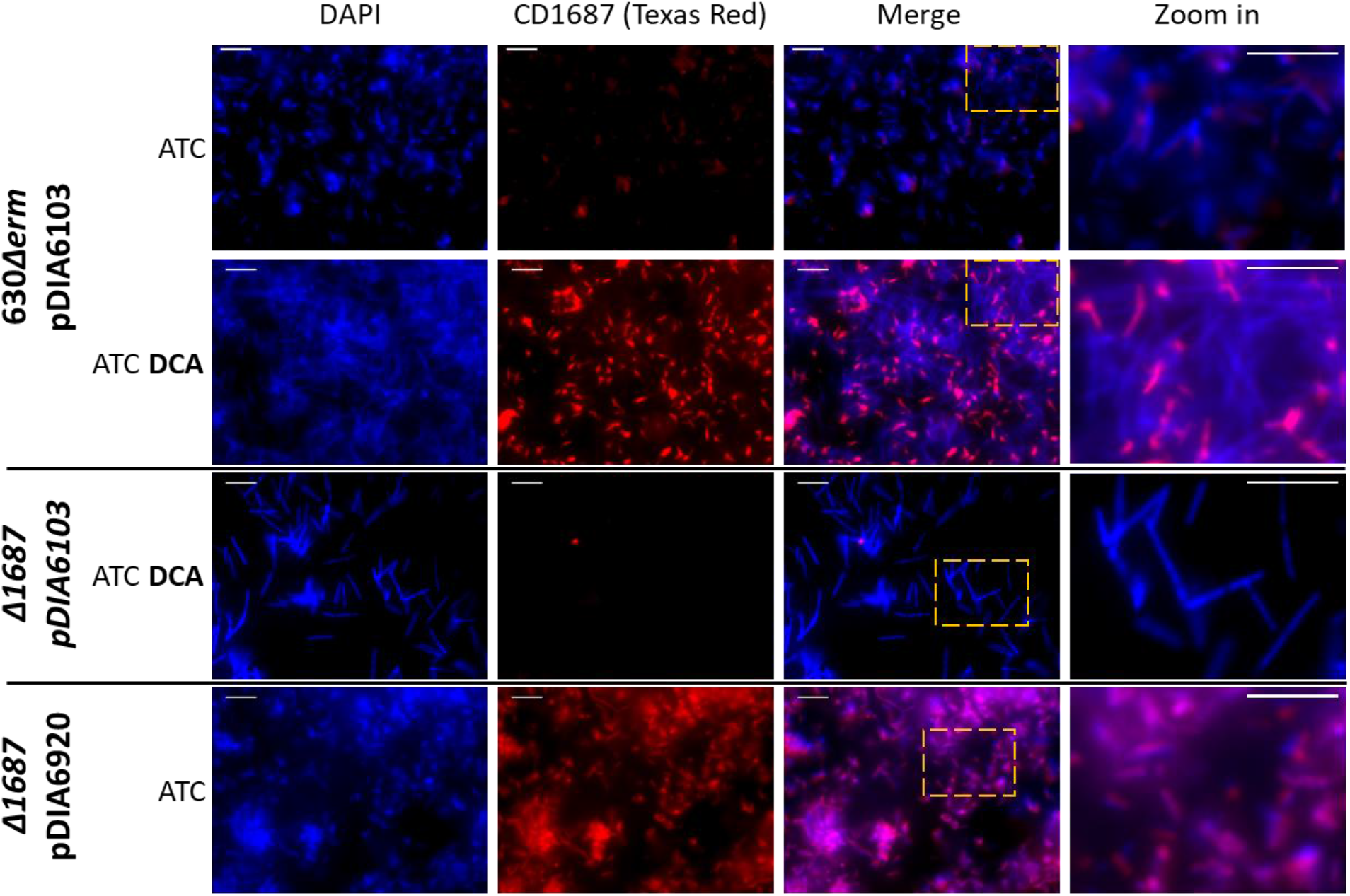
CD1687 localizes at the cell surface of *C. difficile* and displays heterogenous distribution within the biofilm. *In situ* epifluorescence microscopy analysis was performed on 48h biofilms grown in BHISG + ATC (100ng/mL) either in the presence or absence of DCA (240μM) as indicated. The strains tested were the wild type strain (630Δ*erm*) carrying the control vector pDIA6103 and with the *Δ1687* strain carrying the plasmid with an inducible CD1687 (pDIA6920) or the control plasmid (pDIA6103). DNA is stained with DAPI (blue) and CD1687 is labelled with specific anti-CD1687 rabbit antibodies detected with a TexasRed-conjugated goat anti-rabbit antibody (red). Pictures are representative of three biological replicates and were taken with a Nikon Eclipse Ti inverted microscope (Nikon, Japan). Scale bar: 10μm.

Based on the cellular localisation of CD1687, we wondered if the addition of the anti-CD1687 antibodies during growth could prevent DCA-induced biofilm formation. As shown in Fig 5a, the addition of the anti-CD1687 polyclonal antibodies to cells grown under biofilm inducing conditions (BHISG + 240μM DCA) strongly inhibited biofilm formation in a dose-dependent manner. No inhibitory effect was observed when an unpublished non-specific antibody was used at the highest concentration of anti-CD1687 that inhibited biofilm formation (data not shown). Additionally, bacterial growth was unaffected by the antibodies, regardless of the concentration used in the biofilm assays (Fig 5b). Therefore, inhibiting extracellular function of CD1687 prevents biofilm formation, suggesting that the presence of the CD1687 at the surface of the cell wall is critical for DCA-induced biofilm formation.

**Figure 5:**
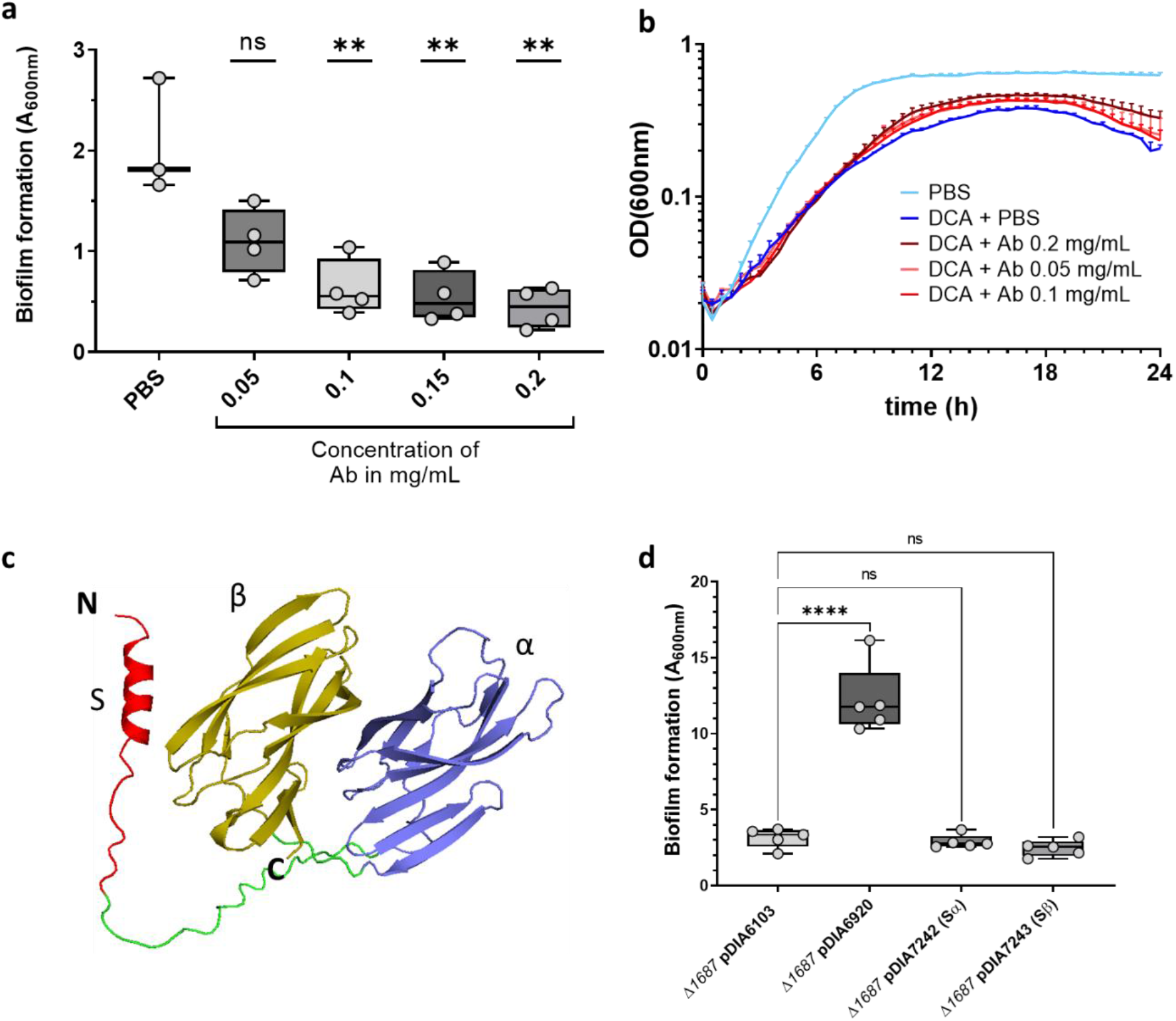
DCA-induced biofilm formation is inhibited in the presence of anti-CD1687 antibodies. **a**. Biofilm formation of the 630*Δerm* strain was assayed 48h in BHISG with DCA (240μM) cultures in presence of different concentration of anti-CD1687 rabbit antibodies (0.05mg/mL to 0.2mg/mL). **b**. Growth kinetics (OD_600nm_) of the WT (630*Δerm*) in BHISG medium with PBS or DCA supplemented with different concentrations of anti-CD1687 rabbit antibodies (0.05mg/mL to 0.2mg/mL). Ab: antibody; nsAb: non-specific antibody. **c**. The alphafold2 predicted structure of CD1687 show a N-terminal signal peptide S (red) connected to the α beta domain (purple) by a linker peptide (green), with another similar β beta domain (yellow) in the C-terminal region. **d**. 48h biofilms form by various *Δ1687* strain complemented with an empty vector (pDIA6103) or plasmids overexpressing the full length CD1687 (pDIA6920) or truncated CD1687 lacking either one of the two domains removed (pDIA7242 and pDIA7243, Table S1) grown in BHISG with ATC (100ng/mL) and DCA (240μM). Each data point represents an independent biological replicate composed of 2 to 4 technical replicates. Asterisks indicate statistical significance with a one-way ANOVA test followed by Dunnett’s multiple comparison test (a) (ns: not significant;**: p<0.01) or a Tukey’s multiple comparison test (d) (ns: not significant;****: p<0.001).

To get some insights on the structure-function of CD1687, we used the software AlphaFold2 (32) to predict the 3D protein structure of CD1687. As shown in Fig 5c, CD1687 has an alpha helix N-terminal signal peptide and two putative beta domains. To search for possible functions of the beta barrels, the putative structure of CD1687 was analysed in the Ekhidna database through the Dali server (33), but no function was detected. Since the function of CD1687 could be assigned to one of the two beta domains, we complemented the *Δ1687* mutant by overexpressing CD1687 with either one of the two domains removed and growing these strains under biofilm inducing conditions (BHISG + 240μM DCA. Complementation of the mutant was not observed, indicating that *C. difficile* needs both beta domainds of the CD1687 to form DCA-induced biofilms (Fig 5d).

### CD1687 binds to DNA in a non-specific manner

Since we did not identify a potential function from the CD1687 structure, we sought to determine if CD1687 has a DNA-binding activity as observed for *Staphylococcus aureus* lipoproteins that promote eDNA-dependent biofilm formation (34). Since the *C. difficile* biofilm matrix is mainly composed of eDNA (15), we tested the ability of CD1687 to bind to DNA by performing an electromobility shift assay (EMSA). When the purified CD1687 protein was incubated with the *E. coli* DNA plasmid pUC9 or a PCR-generated amplicon produced from *C. difficile* DNA (from a sequence in the region of *CD1438*), we observed that the migration of the DNA was shifted by the presence of the CD1687 and increasing CD1687 concentration correlates with more retention (Fig 6ab). However, we did not observe a shift when CD1687 is heat-inactivated or if BSA was used as control at the highest concentration of CD1687 that shift DNA fragments. Therefore, CD1687 can bind DNA and could potentially interact with eDNA when exposed at the surface.

**Figure 6:**
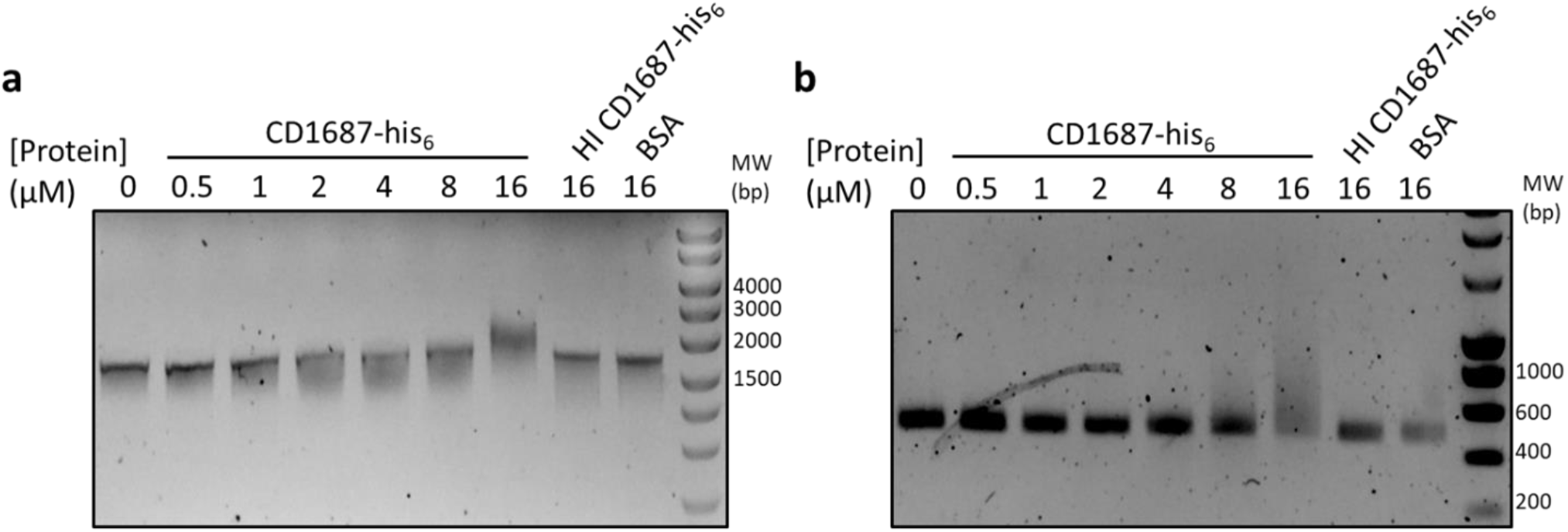
CD1687 binds DNA and shifts DNA migration. Electrophoretic Mobility shift assay (EMSA) was performed with **a**. *E. coli* plasmid pUC9 or **b**. *C. difficile* DNA (450bp PCR-amplicon) mixed with various concentrations of CD1687 (up to 16μM), with 16μM of heat inactivated (HI) CD1687 or BSA used as controls.

## Discussion

In this study, we confirmed that only CD1687 in the *CD1685-CD1689* cluster was required for DCA-induced biofilm formation and this required the localization of CD1687 at the cell surface. Despite this importance, there is a significant heterogeneity in response to DCA for the expression and localization of CD1687 at cell surface in the population as observed by microscopy (Fig 4 and Fig S5). This would explain the relatively low transcriptional level of the *CD1685-CD1689* gene cluster at the population level (15). Interestingly, the more CD1687 is homogeneously expressed in the cell population, the greater the biofilm formed (Fig 4, Fig S2c). To our knowledge, expression heterogeneity of critical biofilm components has not yet been reported in *C. difficile*. Phenotypic heterogeneity in biofilms is well characterized in several other bacterial species resulting in phenotypic diversification and division of labor in a clonal bacterial population (35). For example, a subpopulation of cells synthesize the exopolysaccharides matrix during biofilm formation in *B. subtilis* (36). Phenotypic heterogeneity has been described in planktonic cells of *C. difficile* and this affected the expression of the flagellum and toxins (37). In this case, heterogeneity is controlled by a specific DNA recombination event mediated by RecV (38) and the Rho factor (39). In addition, *C. difficile* colony morphology is also subjected to phenotypic heterogeneity resulting in changes in bacterial physiology and pathogenesis and this occurs through phase variation of the CmrRST signal transduction system expression (40,41).

Given that CD1687 forms an operon with a two-component regulatory system (CD1688-1689) and that CD1687 is a cell wall protein, we first hypothesized that CD1687 was involved in signal transduction leading to transcriptional modifications in response of DCA. However, CD1687 did not bind DCA, which eliminates the role of CD1687 as a DCA sensing protein. Furthermore, we compared genes regulated by CD1688 (42) and, with the exception of sporulation, found limited overlap suggesting that CD1687 may not be part of the CD1688-CD1689 signaling cascade. This is consistent with the absence of CD1689 and CD1688 in our pull-down assay. However, several solute-binding proteins and transporter-associated proteins were isolated in a pull-down assay. This and the transcriptional analysis provide evidence that CD1687 influences the metabolism of *C. difficile*. In support of this, regulators (Spo0A, CodY and SinRR’) that manage metabolic priorities during growth phases, were differentially regulated when CD1687 was overexpressed (26,43,44). Furthermore, the expression of the gene encoding toxin and those involved in sporulation were also affected and these processes are known to be dependent on the metabolic state of *C. difficile*. When we compared the genes differentially regulated in the absence of CD1687 under DCA-inducing conditions to those differentially regulated when CD1687 was overexpressed in the absence of DCA, there were only 69 common genes, which included genes involved in different metabolic pathways and transport. However, these changes in metabolism-associated genes did not overlap with our previous analyses on gene expression during DCA-induced biofilm formation (23), suggesting that CD1687 is not part of the immediate response to DCA and probably plays a role in the downstream response. Taken together, our data suggest that CD1687 helps reorganize metabolic priorities in response to DCA but this hypothesis alone does not explain the role of CD1687 in the biofilm formation without DCA. Therefore, CD1687 may have additional roles.

Interestingly, many proteins found at the bacterial cell surface interact with eDNA found in the biofilm matrix and this contributes to the organization and structural stability of the biofilm (45). Membrane lipoproteins have already been shown to directly interact with eDNA and participate in biofilm architecture. In *S. aureus*, several membrane-attached lipoproteins interacting with the eDNA of the biofilm matrix have been identified as promoting *S. aureus* biofilm formation (34). We demonstrated that CD1687 interacts with DNA in a non-specific manner supporting the hypothesis that CD1687 acts as an eDNA binding protein during biofilm formation by creating anchor points for eDNA on the cell surface. Similar to our observation with CD1687, overexpressing eDNA-binding proteins in *S. aureus* resulted in an increased retention of surface eDNA and an enhanced biofilm biomass. However, deleting the *S. aureus* lipoproteins had minimal impact on biofilm formation but biofilm porosity increase indicating that interactions of the lipoprotein with eDNA contribute to overall biofilm structure. Unlike the lipoprotein found in *S. aureus*, a deletion or inactivation of *CD1687* abolished biofilm formation (34). CD1687 interacting with eDNA seems to be an essential part of DCA-induced biofilm formation. Other structures may also interact with eDNA. Recently, two minor subunits (PilW and PilJ) of the *C. difficile* T4P were shown to directly interact with eDNA to promote biofilm formation (46). Neither subunit have a predicted DNA-binding motif as observed with CD1687. The T4P is a structure that promotes biofilm formation in the absence (47,48) or presence of DCA (23). In the presence of DCA, PilW is upregulated but is not required for biofilm formation (15,23). Furthermore, the *pilW* gene was differently regulated in our transcriptome; up-regulated in the WT vs *Δ1687* with DCA analysis (significantly but below the threshold) and down-regulated in the overexpressed CD1687 vs WT without DCA analysis. Therefore, CD1687 and the T4P may have complementary role and the lack of eDNA-binding by one of these components may change the behavior of *C. difficile* during biofilm formation.

Despite the potential role of CD1687 as an eDNA-binding protein and in metabolism, we cannot exclude that the overexpression of CD1687 modifies the properties of the cell wall through the interactions of CD1687 with other membrane proteins and transporters (Table S5). These interactions could be detected by different sensors, which would activate a feedback loop to modify the cell wall and the composition of the cell surface proteins. For example, the *dltABCD* operon was up-regulated when CD1687 was overexpressed in the absence of DCA. The

DltABCD proteins are responsible for the D-alanylation of teichoic acids, which changes the electrical charges of the cell wall and surface (49). Overexpression of CD1687 also affected cell morphology; in response to DCA, cells expressing high levels of CD1687 show reduced size and shape distortion (Fig 4 and Fig S5). Overall, the overexpression of CD1687 may have downstream effects on the physiology of *C. difficile* and these changes may contribute to biofilm formation.

Finally, our hypothesis is that the mechanism for biofilm formation in the presence of DCA is different than the mechanism when DCA is absent and CD1687 is overexpressed. In the presence of DCA, we know that *C. difficile* goes through a metabolic re-organization (23) and, based on our data, CD1687 would help with metabolic priorities for long term adaptation. Once there is enough eDNA, CD1687 would interact with eDNA binding and serve as an anchor point. When CD1687 is overexpressed independently of DCA, it increases homogeneity of CD1687 surface localization in the population and serves as multiple anchoring sites for eDNA resulting in a strongly adherent biofilm. As observed in *S. aureus*, other lipoproteins may bind eDNA in *C. difficile* and several are upregulated in response to DCA (23). Unlike the lipoproteins characterized in *S. aureus*, the lipoprotein CD1687 probably has a critical function in metabolism in response to DCA and other lipoproteins do not provide functional redundancy. This highlights the importance of CD1687 in promoting biofilm formation. More research will be needed to understand the role and the contribution of these other lipoproteins to biofilm.

## Acknowledgements

This work was funded by the Institut Pasteur, the “Integrative Biology of Emerging Infectious Diseases” (LabEX IBEID) funded in the framework of the French Government’s “Programme Investissements d’Avenir” and The ANR DifBioRel AAPCE5. EA is a doctoral fellow of Université Paris-Cité. LH is a doctoral fellow of Sorbonne Université.

