## Supplementary material for "The cell wall lipoprotein CD1687 acts as a DNA binding protein during deoxycholate-induced biofilm formation in *Clostridioides difficile*"

#### List of contents:

**Supplementary Material and Methods**

**Figure S1:** Effect of DCA on the transcript levels of the *CD1685-CD1689* gene cluster
**Figure S2:** Confirmation of gene deletion in the *CD1685-CD1689* cluster and biofilm formation by the resulting deletion strains.
**Figure S3:** Analysis of the interactions between DCA and immobilized CD1687 by surface plasmon resonance.
**Figure S4:** Functional classification of the differentially expressed genes identified in the transcriptomic analyses.
**Figure S5:** Protein levels and localization of CD1687 on the surface of biofilm grown *C. difficile* cells by immunofluorescence.
**Figure S6:** SDS-PAGE and Western immunoblot of proteins isolated from *E. coli* expressing CD1687 and Western immunoblot of CD1687 protein levels in different *C. difficile* strains.

**Table S1:** Lists of strains, plasmids and primers used in this study
**Table S2:** Sequencing results from the DNA amplicons generated by 5'RACE
**Table S3:** Major transcriptomic changes observed in the transcriptomic experiments
**Table S4:** List of common genes differently expressed in both transcriptomic experiments
**Table S5:** Protein isolated by pull-down

### Supplementary Material and Methods

**Production and purification of CD1687**. *E. coli* strain Bli5 containing a pET20-derived plasmid carrying the *CD1687* gene (Table S1) was used to overexpress hexa-histidine-tagged CD1687 protein without its signal peptide. Cells were grown overnight at 37°C in LB supplemented with glucose (1% w/v) and antibiotics (ampicillin 100µg/mL and chloramphenicol 15µg/mL). The overnight culture was transferred (1/100) in 1 L of the same medium and incubated at 37°C. Once the culture reached an $OD_{600nm}$ of 0.5, IPTG was added (final concentration 0.1mM) and the culture was incubated for an additional 3h. Cells were then harvested by centrifugation (5000g, 10min, 4°C) and the pellet was washed with cold PBS. After centrifugation, the supernatant was discarded and resulting pellet was frozen at -20°C. The pellet was then resuspended in 15 mL of lysis buffer (50mM sodium phosphate pH=8.0; 300mM NaCl) and sonicated. After centrifugation (5000g, 10min, 4°C), the supernatant was collected and mixed with Ni-NTA beads and incubated one hour at 4°C. The beads were then transferred to an elution column and washed with washing buffer (50mM sodium phosphate pH=8.0; 300mM NaCl; 10mM imidazole). Proteins were eluted with 2 ml of sodium phosphate buffer (50mM, pH=8.0) supplemented with 300mM NaCl and a gradient of imidazole ranging from 50mM to 500mM. Eluted proteins were analysed by western immunoblotting and fractions containing CD1687 were dialyzed in TAE buffer (Tris-base (20mM); acetic acid (10mM); EDTA (0,5mM); pH = 8.5) using Slide-A-Lyzer dialysis units (Thermo Fisher Scientific, USA). To raise polyclonal anti-CD1687 antibodies, two female rabbits (New Zealand White) were injected four times with 50 µg of purified CD1687($His_6$) (0.5mL of antigen with 0.5mL of complete Freund's adjuvant at D0, D14, D28 and D42) with the Covalab company (France). Antibodies were purified at D53 of immunization.

**Real-time surface plasmon resonance binding assay.** All experiments were performed on a Biacore T200 instrument (Cytiva, USA) equilibrated at 25°C in buffer TAE (20mM Tris base, acetic acid 10mM, EDTA 0.5mM, pH = 8.5). CD1687($His_6$) (100µg/ml) was captured for 600s at 2µl/min on an NiCl2-loaded NTA sensorchip, reaching a surface density of 1000-1200 RU (resonance units; 1RU ≈1pg/mm$^2$). DCA (16-2000µM) was then injected at 10µl/min for 120s, simultaneously on the CD1687 surface and on an empty reference chip from which non-specific signals were subtracted.

**Protein sequencing assay via mass spectrometry.**

**Protein Digestion.** Proteins were reduced using 5mM TCEP for 30min at room temperature. Alkylation of the reduced disulfide bridges was performed using 10mM iodoacetamide for 30min at room temperature in the dark. Proteins were then digested in two steps, first with 250 ng r-LysC Mass Spec Grade (Promega) for 4h at 30°C then samples were diluted below 2M urea with 100mM Tris HCl pH 8.5 and 500ng Sequencing Grade Modified Trypsin was added for the second digestion overnight at 37°C. Proteolysis was stopped by adding formic acid (FA) at a final concentration of 5%. The resulting peptides were cleaned using AssayMAP C18 cartridges on the AssayMAP Bravo platform (Agilent) according to the manufacturer's instructions. Peptides were concentrated to dryness and resuspended in 2% acetonitrile (ACN) and 0.1% FA just prior to LC-MS injection.

**LC-MS/MS analysis.** LC-MS/MS analysis was performed on a Q ExactiveTM Plus Mass Spectrometer (Thermo Fisher Scientific) coupled with a Proxeon EASY-nLC 1200 (Thermo Fisher Scientific). 500ng of peptides were injected onto a home-made 37cm C18 column (1.9µm particles, 100Å pore size, ReproSil-Pur Basic C18, Dr. Maisch GmbH, Ammerbuch-Entringen, Germany). Column equilibration and peptide loading were done at 900 bars in buffer A (0.1% FA). Peptides were separated with a multi-step gradient from 3 to 6% buffer B (80% ACN, 0.1% FA) in 5 min, 6 to 31 % buffer B in 80min, 31 to 62% buffer B in 20min at a flow rate of 250nL/min. Column temperature was set to 60°C. MS data were acquired using Xcalibur software using a data-dependent method. MS scans were acquired at a resolution of 70,000 and MS/MS scans (fixed first mass 100m/z) at a resolution of 17,500. The AGC target and maximum injection time for the survey scans and the MS/MS scans were set to 3E6, 20ms and 1E6, 60ms respectively. An automatic selection of the 10 most intense precursor ions was activated (Top 10) with a 30s dynamic exclusion. The isolation window was set to 1.6m/z and normalized collision energy fixed to 27 for HCD fragmentation. We used an underfill ratio of 1.0% corresponding to an intensity threshold of 1.7E5. Unassigned precursor ion charge states as well as 1, 7, 8 and >8 charged states were rejected and peptide match was disable.

**Protein identification and quantification.** Acquired Raw data were analyzed using MaxQuant software version 2.1.1.0 (1) using the Andromeda search engine (2,3). The MS/MS spectra were searched against the Clostridium difficile 630 database (3,957 entries).

All searches were performed with oxidation of methionine and protein N-terminal acetylation as variable modifications and cysteine carbamidomethylation as fixed modification. Trypsin was selected as protease allowing for up to two missed cleavages. The minimum peptide length was set to 7 amino acids and the peptide mass was limited to a maximum of 4,600Da. The false discovery rate (FDR) for peptide and protein identification was set to 0.01. The main search peptide tolerance was set to 4.5ppm and to 20ppm for the MS/MS match tolerance. Second peptides was enabled to identify co-fragmentation events. A false discovery rate cut-off of 1% was applied at the peptide and protein levels. The mass spectrometry proteomics data have been deposited to the ProteomeXchange Consortium via the PRIDE partner repository with the dataset identifier PXD038282. The statistical analysis of the proteomics data was performed as described previously (4). Briefly, four biological replicates were acquired per condition. To highlight significantly differentially abundant proteins between two conditions, differential analyses were conducted through the following data analysis pipeline: (1) deleting the reverse and potential contaminant proteins; (2) keeping only proteins with at least two quantified values in one of the two compared conditions to limit misidentifications and ensure a minimum of replicability; (3) log2-transformation of the remaining intensities of proteins; (4) normalizing the intensities by median centering within conditions thanks to the normalizeD function of the R package DAPAR (5), (5) putting aside proteins without any value in one of both compared conditions: as they are quantitatively present in a condition and absent in another, they are considered as differentially abundant proteins and (6) performing statistical differential analysis on them by requiring a minimum fold-change of 2 between conditions and by using a LIMMA t test (6,7) combined with an adaptive Benjamini-Hochberg correction of the p values thanks to the adjust.p function of the R package cp4p (8). The robust method of Pounds and Cheng was used to estimate the proportion of true null hypotheses among the set of statistical tests (9). The proteins associated with an adjusted p value inferior to an FDR level of 1% have been considered as significantly differentially abundant proteins. Finally, the proteins of interest are therefore the proteins that emerge from this statistical analysis supplemented by those being quantitatively absent from one condition and present in another.

### Supplementary figures

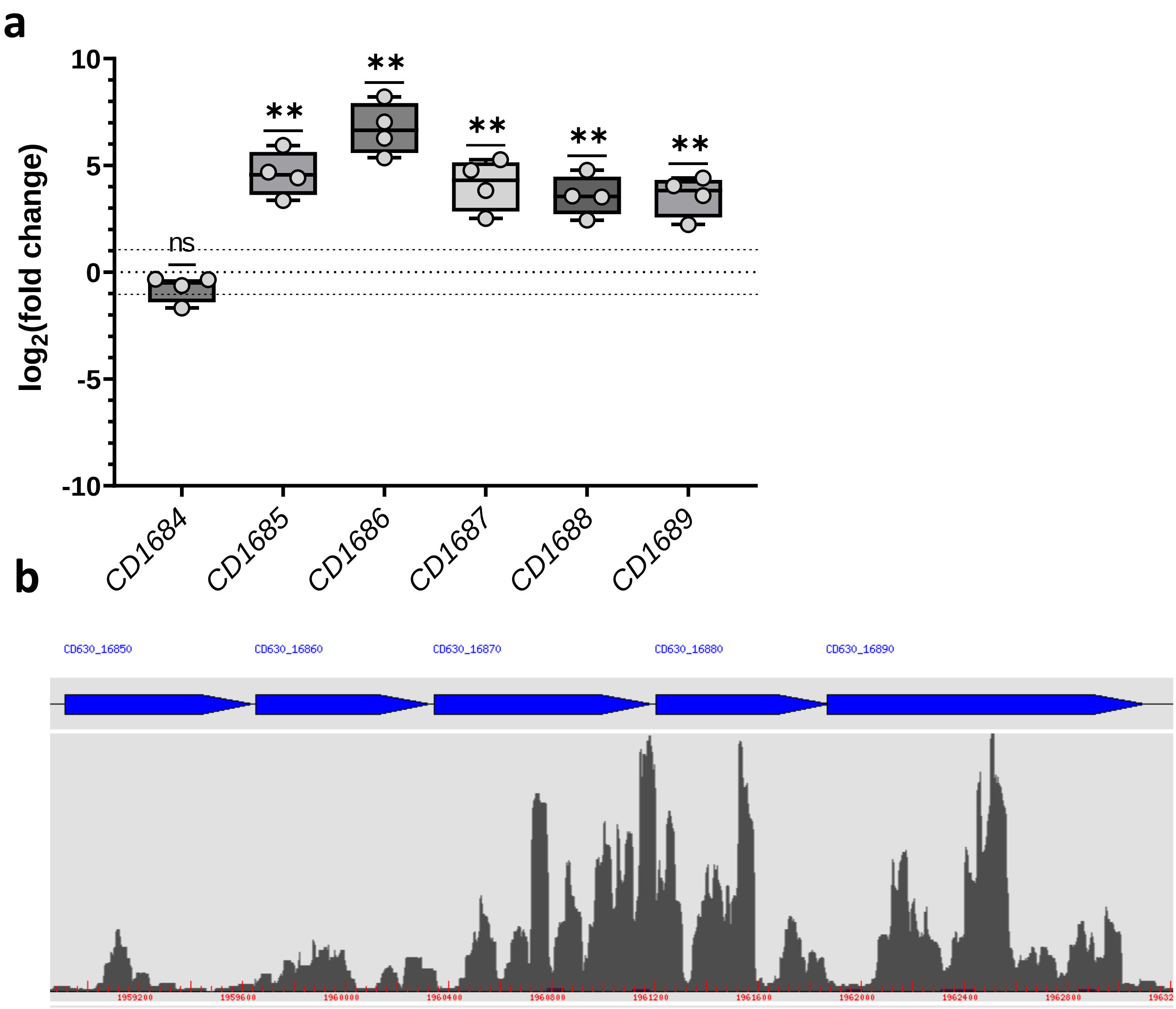


**Figure S1: Effect of DCA on the transcript levels of the *CD1685-CD1689* gene cluster. a.** RNA was extracted from *C. difficile* strain 630*Δerm* grown for 48h in BHISG with or without DCA (240µM). Comparative RT-qPCR analysis of the *CD1685-CD1689* genes were performed between cell grown in the presence and in the absence of DCA. **b.** Raw read numbers detected along the *CD1685-CD1689* region provided by the transcriptome published in Dubois *et al.* (2019) from cells grown in BHISG with 240µM DCA for 48h.
Asterisks indicate statistical significance with a t test comparing the theoretical mean of 0 (ns: not significant; **: p<0.01)

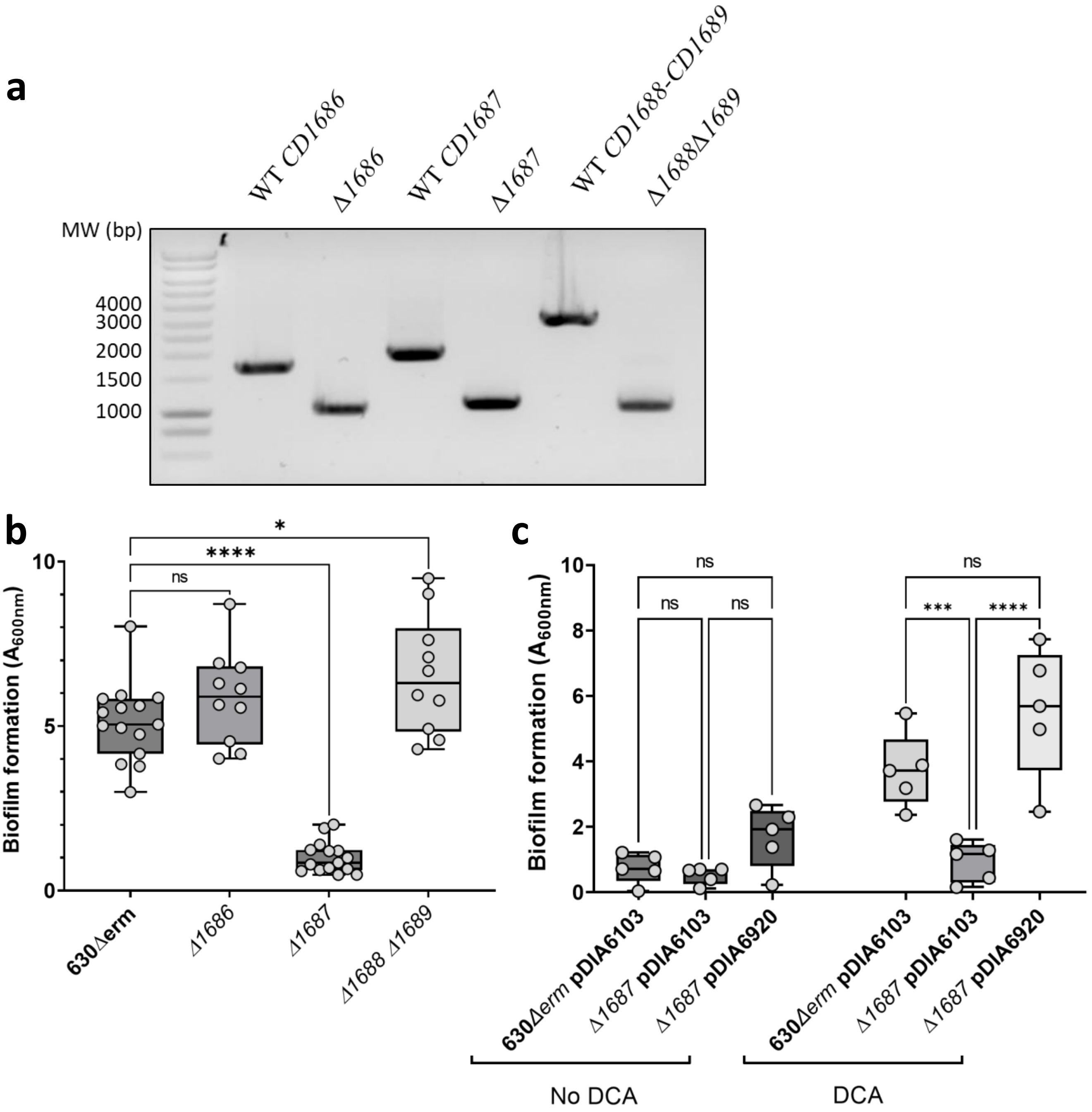


**Figure S2: Confirmation of gene deletion in the *CD1685-CD1689* cluster and biofilm formation by the resulting deletion strains. a** PCR amplification using DNA from the wild type strain, the *Δ1686 strain*, *Δ1687* strain and the *Δ1688Δ1689* strain was performed using the "verification" primers listed in Table S1. In the WT/630*Δerm* strain, the expected sizes of the PCR products were 1625bp, 1820bp, and 2809bp for *CD1686*, *CD1687* and *CD1688-CD1689* genes, respectively. In the deletion strains, the expected sizes of the PCR products were 968bp, 995bp and 939bp for the *Δ1686, Δ1687,* and *Δ1688- Δ1689* deleted genes, respectively. **b.** Biofilms formation by the wild type (630*Δerm*), the *Δ1686,* the *Δ1687* and the *Δ1688Δ1689* strains was assayed 48h after inoculation in BHISG with DCA (240µM), **c.** Biofilms formation by wild type strain complemented with the control plasmid (pDIA6103) and the *Δ1687* mutant strain complemented with the *CD1687* plasmid (pDIA6920) or the control plasmid (pDIA6103) was assayed 48h after inoculation in BHISG with ATC (100ng/mL), in the presence or not of DCA (240µM). Each data point represents an independent biological replicate composed of 2 to 4 technical replicates.
Asterisks indicate statistical significance with a one-way ANOVA test followed by a Tukey's multiple comparison test (ns: not significant; *: p<0.05; ***: p<0.001; ****: p<0.0001).

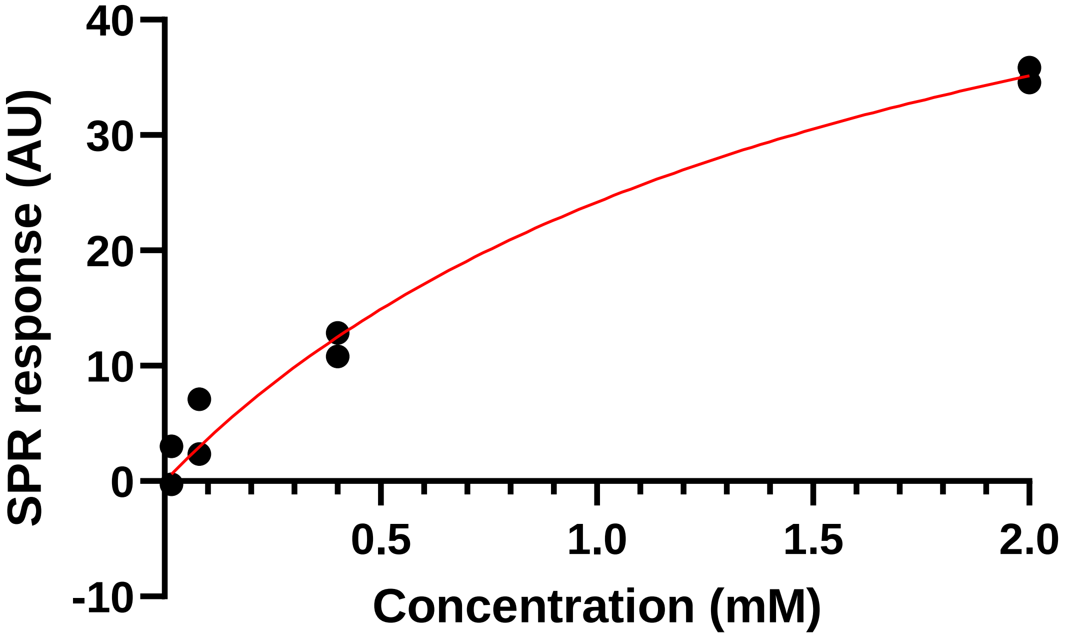


**Figure S3:** Analysis of the interactions between DCA and immobilized CD1687 by surface plasmon resonance. The specific steady-state surface plasmon resonance responses were determined and plotted against the DCA concentration, allowing to determine the affinity and stoichiometry of the interaction between CD1687 and DCA. Dots represent the raw data while the red curve represents the fitted model. Here we determined that the dissociation constant of the DCA-CD1687 complex is 1.65±0.58mM and the stoichiometry is of 5±1 DCA molecule per CD1687 protein.

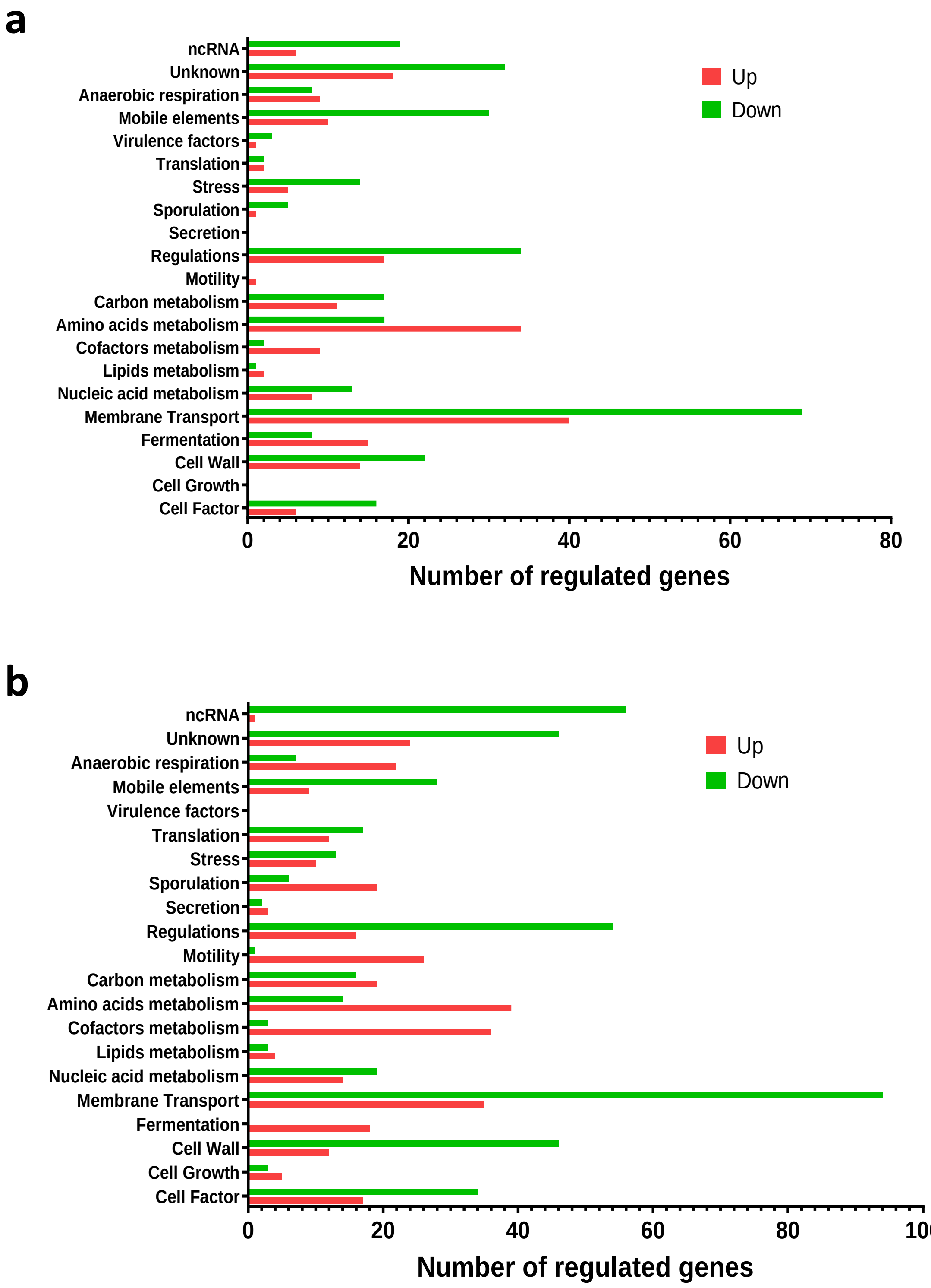


**Figure S4**: **Functional classification of the differentially expressed genes identify in the transcriptomic analyses. a.** Transcriptome comparison between the wild type 630*Δerm* and the Δ*1687* mutant strain grown in BHISG in presence of DCA (240µM) for 24h. **b.** Transcriptome comparison between the wild type strain carrying an inducible plasmid overexpressing CD1687 (pDIA6920) grown in BHISG in the presence and absence of the ATC inducer for 24h. Changes in expression are colour coded (in red, up-regulated; or green, down-regulated). Genes were classified by functional classes and the numbers of genes are reported on the x-axis of the graphs.

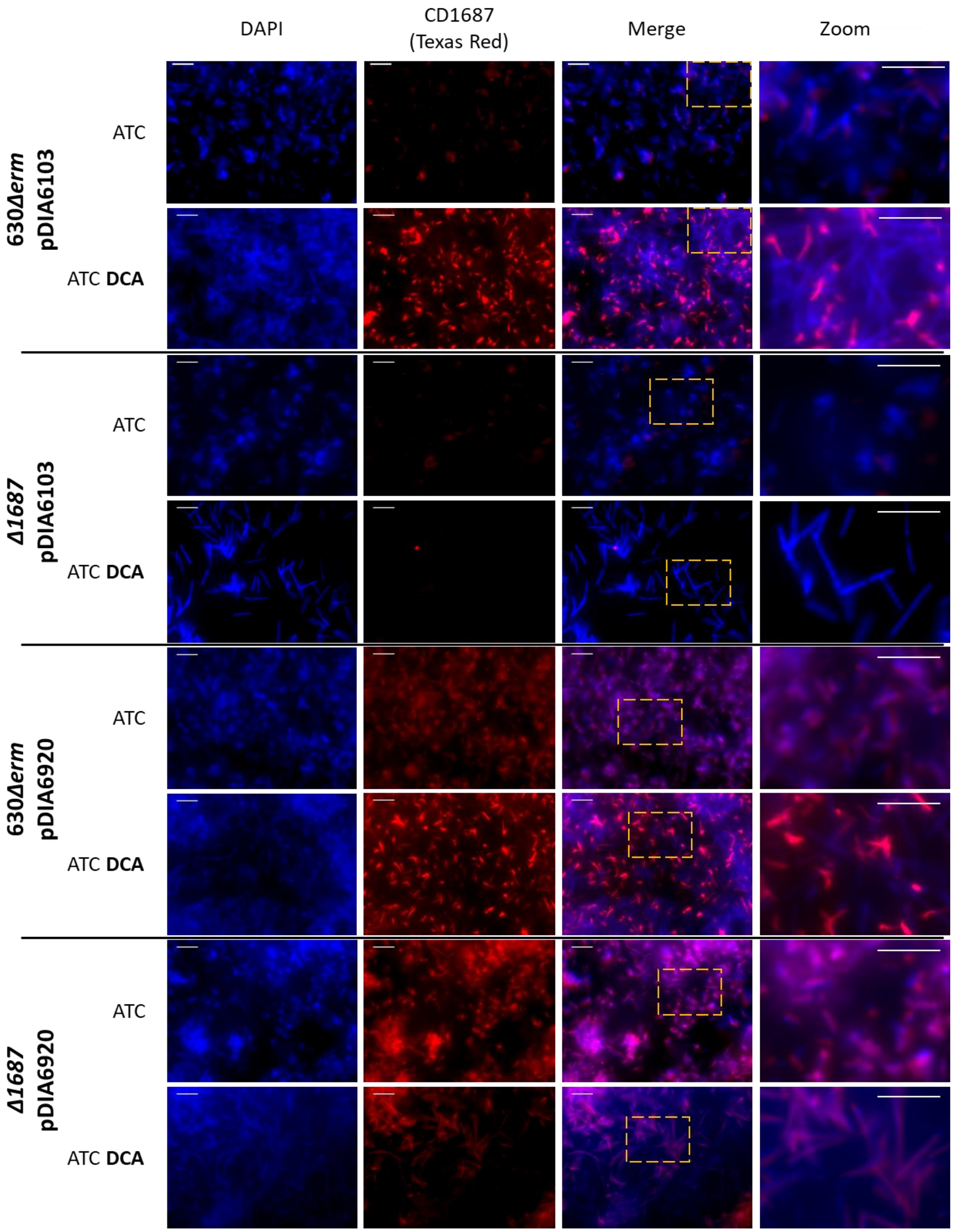


**Figure S5:** Protein levels and localization of CD1687 at the cell surface of *C. difficile* cells within biofilm by immunofluorescence. *In situ* epifluorescence microscopy analysis was performed on the wild type strain (630*Δerm*) and the *Δ1687* mutant strain carrying the *CD1687* plasmid (pDIA6920) or the empty control plasmid (pDIA6103) grown as 48h biofilm in BHISG with ATC (100ng/mL), with or without DCA (240µM). DNA was stained with DAPI (blue) and CD1687 was detected with a specific anti-CD1687 rabbit antibody detected with a Texas Red-conjugated goat anti-rabbit antibody (red). Pictures are representative of three biological replicates and were taken with a Nikon Eclipse Ti inverted microscope (Nikon, Japan). Scale bar: 10µm.

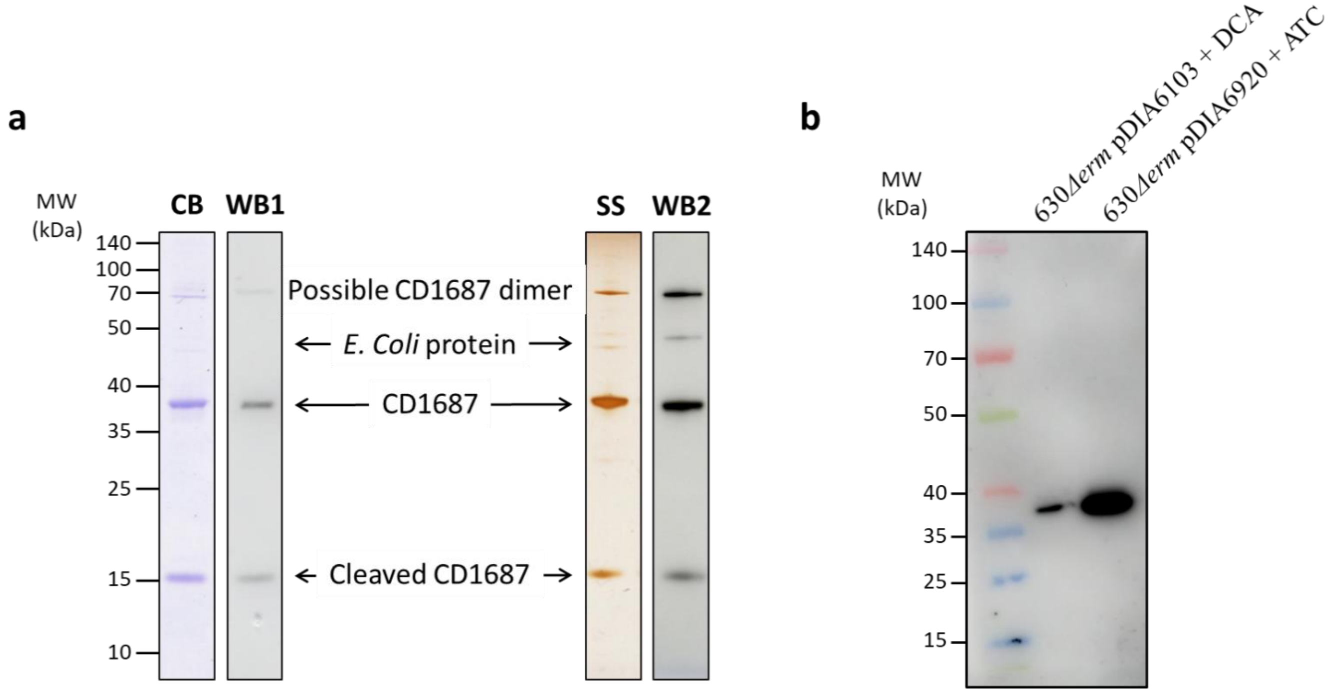


**Figure S6:** SDS-PAGE and western immunoblot of the CD1687 protein expressed from *E. coli* or from different *C. difficile* strains. **a.** Analysis of the protein purified from *E. coli* expressing the CD1687-his$_6$ protein by silver staining (SS), Coomassie blue coloration (CB) and Western Blot (WB) using either a mouse anti-his$_6$ antibody (Invitrogen) (WB1) or an anti-CD1687 rabbit antibody (WB2). The CD1687 dimer and the CD1687 truncated protein form of CD1687 were detected only in *E. coli* extracts. **b.** CD1687 protein levels detected by western immunoblot using an anti-CD1687 antibody from protein extracted from the wild type with the control vector pDIA6103 or the complementation *CD1687* plasmid (pDIA6920). The bacteria were grown for 48h in BHISG supplemented either with ATC (100ng/mL) or DCA (240µM). 2µg of total proteins were loaded in each running lane.

### Supplementary Tables

**Table S2:** Sequencing results from the DNA amplicons generated by 5'RACE

| | Last sequenced 30bp | TSS | 50bp upstream |
|---|---|---|---|
| 1 | GAATATATGATGTCTATACAAGATATAAAA | TSS1 | TGATGTTTGATAGGGGAAAATGGAATAAATTGGATTATAATGAACTTTTA |
| 2 | GGAAATTGGAAGGAATTTTTAGAAGTAGCA | TSS1 | TAGGTATTAGAGTTCCTAATTTAAGAAAGTTATCAAAAGAAATAAGCCAA |
| 3 | ACATATTAAAGAAATATTTGAATACTGTAA | TSS2 | GAAATTAGATTTGCATTAGTTATGTTTTTAATATATTATGTTGATGATAA |
| 4 | ACATATTAAAAGACGAATTAAATTATAATA | TSS3 | TTTAAGAACTTTTTTAAATAAAAAAGGATATAAGTCAGAAAAAGGTAAAT |
| 5 | CATATTAAAAGACGAATTAAATTATAATAA | TSS3 | TTAAGAACTTTTTTAAATAAAAAAGGATATAAGTCAGAAAAAGGTAAATA |
| 6 | ATTAAATTATAATAAAAGTTTGGATAATAG | TSS3 | AATAAAAAAGGATATAAGTCAGAAAAAGGTAAATACATATTAAAAGACGA |
| 7 | ACATATAGATGATGAACACATAGATGAATT | TSSa | GTAAAAGATTACAATATAAAGCCTAAAAAAATATCTAATAAAAAAGATGT |
| 8 | ACACATAGATGAATTAAAAACAACAAGAAT | TSSa | ATAAAGCCTAAAAAAATATCTAATAAAAAAGATGTACATATAGATGATGA |
| 9 | TATAAAAATCTACTATAAAATACTACTATA | TSSb | TTGAAAAACATCAAATACTTACACAGGCATTGAAAGAATTTATGAAAAAG |

red: putative -10 box ; green: putative -35 box
